# Transcriptional and neuroprotective effects of hexokinase-2 inhibitors administered after stroke

**DOI:** 10.1101/2025.04.11.648425

**Authors:** Seok Joon Won, Gӧkhan Uruk, Nguyen Mai, Chia-Ling Tu, Devran Ogut, Sona Asatryan, Ebony Mocanu, Rachel Shon, Kyungsoo Kim, Khukheper Awakoaiye, Kajsa Arkelius, Wenhan Chang, Neel S. Singhal, Raymond A. Swanson

## Abstract

The inflammatory response induced by stroke can exacerbate injury to peri-infarct tissue. Microglia and other immune cells that mediate this response require increased glycolytic flux during pro-inflammatory activation. These cells, unlike neurons and most other cell types, utilize hexokinase-2 (HK2) rather than hexokinase-1 for glycolysis, such that HK2 inhibitors may selectively target them to suppress post-ischemic inflammation. Here we compared the effects of the non-selective hexokinase inhibitor 2-deoxyglucose to the HK2-selective inhibitors lonidamine and 3-bromopyruvate on secondary injury after stroke. A spatial transcriptomic assessment was performed in parallel to compare effects of the inhibitors on microglial gene expression and microglia - neuron interactions and to screen for off-target effects. Each of the inhibitors suppressed pro-inflammatory gene upregulation in peri-infarct microglia and attenuated the upregulation of cell stress functional pathways in the neighboring neurons, but had minimal effect on neuronal gene expression in uninjured cortex. The HK2-selective inhibitors were more effective than 2-deoxyglucose in suppressing morphological microglial changes, neuronal oxidative stress, and neurite loss. 3-bromopyruvate administered after stroke produced long-term improvements in functional outcome. Selective HK2 inhibitors may thus provide a clinically applicable means to suppress microglial activation and thereby improve outcomes after stroke without endangering neuronal energy metabolism.

---

Focal ischemic stroke results from occlusion of an artery to the brain or spinal cord, usually by a blood clot, and causes infarction (pan-necrosis) in the vascular territory involved. Injury resulting from stroke can be mitigated by clot lysis or mechanical removal, but only a small minority of stroke patients can be treated safely or effectively by these methods ^1, 2^. However, in addition to the tissue infarction that occurs within hours after stroke, a delayed secondary injury develops in the peri-infarct tissue. Inflammation is a major cause of this secondary injury, and since the post-stroke inflammatory response takes many hours to days to fully develop, anti-inflammatory intervention is recognized as a clinically feasible approach that could potentially benefit almost all stroke patients ^3, 4^.

The post-stroke inflammatory response is triggered by alarmins and other factors diffusing out from the infarct. It consists of an initial microglial activation followed by reactive astrocytosis and influx of macrophages and other immune cell types ^3, 4^. This innate immune response is an evolutionarily conserved first line of defense against microbial infections, and as such it entails release of cytotoxic reactive oxygen species, proteases, and cytokines .

This oro-inflammatory microglial activation requires a large increase in glycolytic flux ^5, 6^. The first step of glycolysis (glucose → glucose-6-phosphate) is catalyzed by hexokinase, of which there are five mammalian isoforms ^7, 8^. Hexokinase-1 is the isoform most abundantly expressed by most cell types in brain, including neurons, astrocytes, oligodendrocytes, and vascular cells, but microglia and other immune lineage cells differ in that they express predominately hexokinase-2 (HK2), and further upregulate HK2 expression when activated ^9, 10^. It remains uncertain why immune cells express predominately HK2 rather than HK1 and require increased glycolysis when activated ^11^, but these properties nevertheless identify HK2 as a target for suppressing post-stroke inflammation.

HK2-selective inhibitors have previously been shown to suppress the innate immune response and improve outcomes in mouse models of sepsis, arthritis, Alzheimer’s disease, and other disorders ^9, 10, 12, 13, 14, 15, 16^. Selective HK2 inhibitors are also in clinical trials for other conditions and have an excellent safety profile ^8, 17, 18^ In contrast to pharmacological HK2 inhibition, genetic HK2 downregulation produces variable and conflicting effects, particularly when the HK2 ablation is complete ^9, 15, 19^. This difference between effects of the pharmacological inhibitors and genetic downregulation may be attributable to loss of protein scaffolding or other functions of HK2, which are distinct from its glucose-phosphorylating function.

The present study aimed to evaluate effects of HK2 inhibition when initiated at a time point with clinical applicability, several hours after stroke. We compared the selective HK2 inhibitors lonidamine (LND) and 3-bromopyruvate (3BP) to each other and to the non-selective hexokinase inhibitor 2-deoxyglucose (2DG) ^20^. Whole transcriptome data were acquired from microglia and neurons in peri-infarct cortex to determine drug effects on microglial gene expression and microglia - neuron interactions, and to screen for off-target drug effects.

Histological assessments performed in parallel were used to correlate microglial gene expression with microglial morphology changes after stroke and to determine HK2 inhibitor effects on secondary neuronal injury in the peri-infarct cortex. These studies were accompanied by a long-term survival study to evaluate effects on motor impairment and recovery after stroke.

## RESULTS

We used the selective HK2 inhibitors lonidamine (LND) and 3-bromopyruvate (3BP) at doses previously shown to inhibit HK2 activity in mouse brain ^9, 15^. These drugs were compared to the non-selective hexokinase inhibitor 2-deoxyglucose (2DG) used at a dose comparable to that previously used in studies of mouse brain ischemia ^22, 23^. 2DG induced a mild sedation, whereas LND and 3BP had no discernible behavioral effect. The inhibitors were administered 3 hours following stroke, and brains were harvested 9 hours later for evaluation of microglial and neuronal gene expression by digital spatial profiling. Whole-transcriptome data were obtained from the peri-ischemic cortex abutting the infarct (denoted “peri-infarct-1”); from the adjacent region spanning 0.5 - 1.0 mm from the infarct edge (“peri-infarct-2”); and from the homologous contralateral uninjured cortex (“contralateral”; Fig. 1). After filtering and quality control, the normalized count data represented 19,740 genes (Supplemental Data Files 1 & 2). Cells identified as neurons or microglia by immunolabeling showed the expected enriched expression of these cell-specific gene sets ^24, 25^. T-distributed stochastic neighbor embedding further confirmed the segregated patterns of gene expression (Fig. 1c,d). Although microglia and macrophages have different embryonic lineages, activated resident microglia, and infiltrating macrophages have very similar phenotypes and gene expression patterns ^21^. For this reason and for brevity, we will not distinguish between these two cell types but instead refer to them collectively as “microglia” in the remainder of this work.

**Figure 1.**
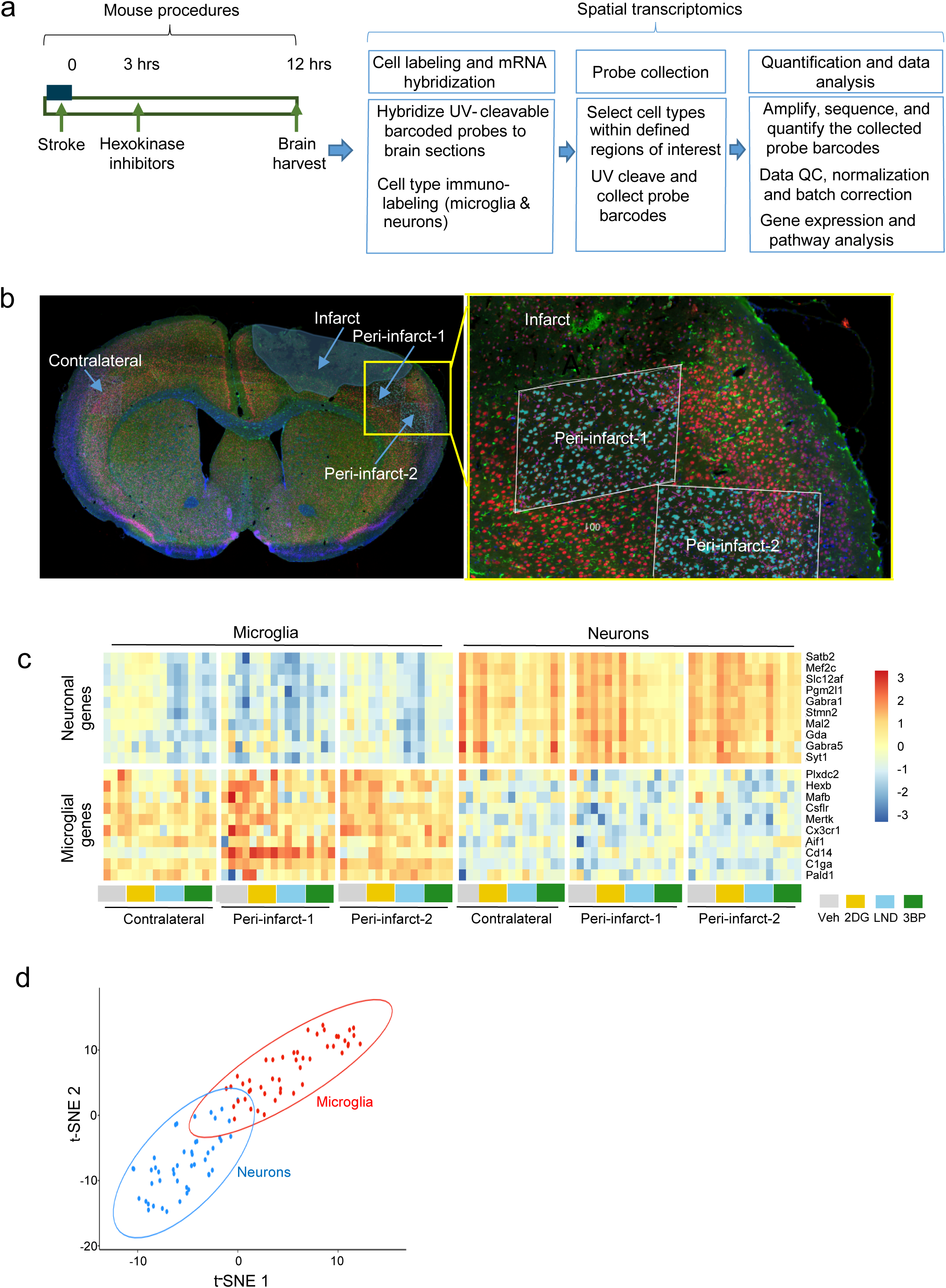
Spatial gene expression analysis of peri-infarct cortex. **a.** Schematic of experimental workflow. **b.** Immunolabeled brain section showing the spatial relationships between the infarct margin and the peri-infarct and contralateral probe collection areas. Microglia are identified by Iba1 immunoreactivity (green) neurons by NeuN (red) and nuclei by DAPI (blue). The yellow box in the low magnification view (left) shows location of the high magnification view (right). In the high magnification view, white parallelograms denote regions of interest in which the DNA barcodes were cleaved from the hybridization probes by UV light. Iba1-labeled cells selected for UV cleavage are displayed here in magenta. **c.** Prototypical neuronal and microglial gene expression patterns corroborate the expected cell identity gene sets in the neurons and microglia irrespective of sampling location or drug treatment group (2DG, 2-deoxyglucose; LND, lonidamine; 3BP, 3-bromopyruvate. (n = 4). **d.** T-distributed stochastic embedding (t-SNE) of normalized expression counts confirms segregation of gene expression patterns by cell type across all treatment conditions. Ellipses represent 95% confidence intervals.

### HK2 inhibitors suppress pro-inflammatory gene expression in peri-infarct microglia

In mice treated with vehicle only, microglia that were close to the infarct edge (the “peri-infarct-1” region) showed a robust increase in pro-inflammatory gene expression relative to microglia in the contralateral cortex. Microglia in the more distant peri-infarct-2 region showed a smaller magnitude but otherwise similar pattern of gene upregulation (Fig. 2a,b). Effects of the HK2 inhibitors were assessed as comparisons between the peri-infarct vs. contralateral cortex under each drug treatment condition (Fig. 2a,b), and as comparisons between each of the HK2 inhibitors vs. vehicle in peri-infarct cortex (Suppl. Fig. 1). By either method, the effects of 3BP and LND were very similar to one another, consistent with a shared mechanism of action. Full data sets for each of the microglial gene expression comparisons are provided as Supplemental Data Files 3 - 7.

**Figure 2.**
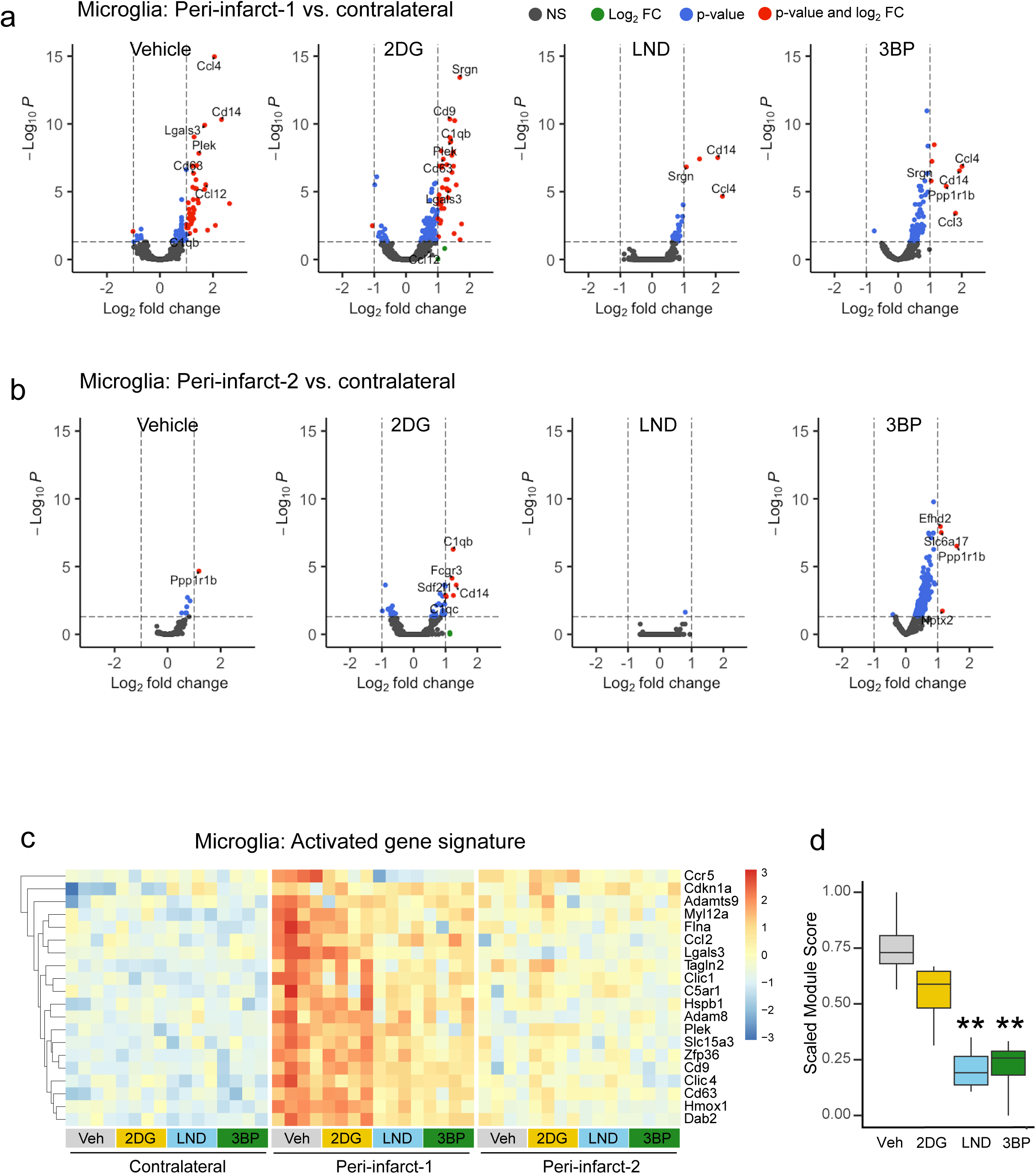
Effects of hexokinase inhibitors on peri-infarct microglial gene expression. Volcano plots show gene expression changes in peri-infarct-1 microglia (**a**) and peri-infact-2 microglia (**b**) relative to microglia in the contralateral cortex under each drug treatment condition (2DG, 2-deoxyglucose; LND, lonidamine; 3BP, 3-bromopyruvate). Genes with log_2_ expression fold change (FC) > 1 and p < 0.05 are denoted by red dots; log_2_ FC > 1 and p > 0.05 by green dots; log_2_ FC < 1 and p < 0.05 denoted by blue dots, and not significant (NS) by black dots. **c**. Heatmap showing relative expression of genes in the ischemia-activated gene set in each drug treatment condition and region of interest n = 4. **d.** Box plot showing the normalized composite expression of the gene sets in peri-infarct-1 cortex. ** p < 0.01 vs. vehicle by Dunnett’s test.

A random forest classification and variable importance analysis were used to identify an activated gene module most characteristic of peri-infarct-1 microglia. Genes in this module included heme-oxygenase 1, the metalloproteinase genes ADAMTS9 and ADAM8, the chemokine receptor CCR5 and ligand CCL2, and others (Fig. 2c). The upregulation of this gene module in peri-infarct cortex was suppressed in mice receiving the HK2 selective inhibitors LND or 3BP, while 2DG had a smaller and non-significant effect as determined with a scaled module score (Fig. 2d, Suppl. Fig. 2). The activated gene module identified here in peri-infarct microglia showed individual gene and pathway overlap with a gene set previously identified as characteristic of disease-activated microglia (DAM) in neurodegenerative diseases ^24, 26, 27^. An evaluation of the DAM gene set showed it to be likewise upregulated in peri-infarct cortex and attenuated by the HK2 inhibitors (Suppl. Fig. 3).

HK2 expression is reportedly higher at baseline in microglia and other myeloid cells than in neurons or other cell types, and further increased by pro-inflammatory activation ^10, 15, 19^. Our results agree with these reports, as we found that microglia in uninjured cortex exhibit a roughly 3-fold higher level of HK2 gene expression than neurons, with this increasing to 12-fold higher in peri-infarct-1 microglia (Fig. 3a). A smaller but statistically non-significant increase was observed in peri-infarct-2 microglia. Surprisingly, these increases in HK2 expression were negated by 3BP or LND and partially negated by 2DG. By contrast, the expression of HK2 in neurons was unaffected by either proximity to the infarct margin or the HK2 inhibitors.

**Figure 3.**
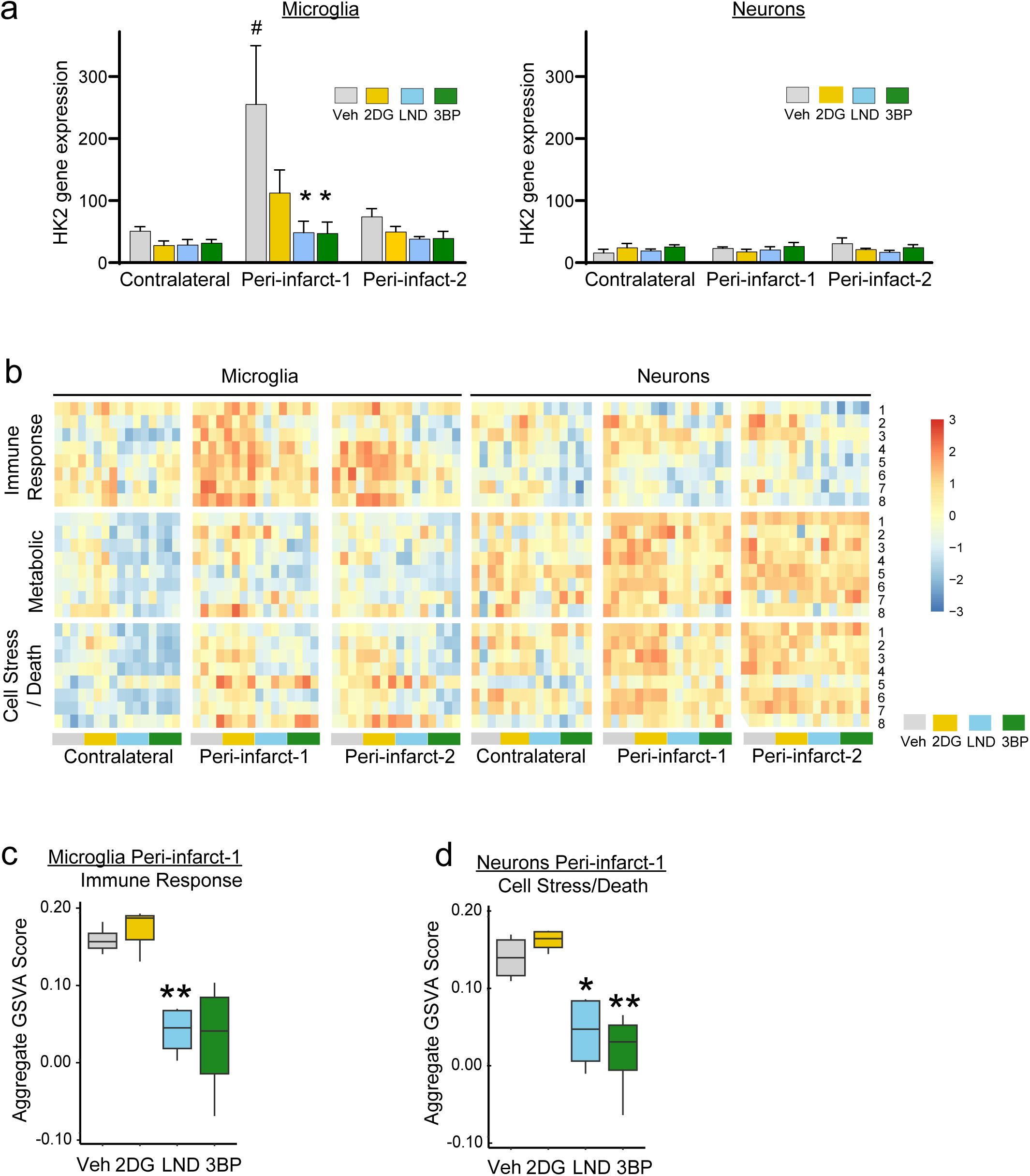
Microglial and neuronal HK2 expression and functional pathway analyses. **a.** HK2 inhibitor effects on HK2 gene expression. n = 4. ^#^ p < 0.01 vs. contralateral; * p < 0.05 vs. vehicle. **b.** Heatmap comparing z-scaled expression of the selected functional gene sets. The functional pathways within each pathway group are denoted by numbers on the right-hand side of the heatmap and listed in Supplemental Figure 5. **c, d.** HK2 inhibitor effects on aggregated gene set variance analysis (GSVA) scores for the microglial immune response functional pathway and the neuronal cell stress/death functional pathway in peri-infarct-1 cortex.

### HK2 inhibitors indirectly affect neuronal gene expression

Although neuronal HK2 expression was not increased in peri-infarct cortex, the whole-transcriptome analysis revealed expression changes in many other neuronal genes (Suppl. Fig. 4, Supplemental Data Files 8 - 12). The HK2-selective inhibitors attenuated these peri-infarct neuronal gene expression changes, while 2DG had less effect. Functional pathways affected by peri-infarct location and the HK2 inhibitors were evaluated by gene set variation analysis (GSVA) using Gene Ontology (GO): Biological Pathway (BP) annotations corresponding to inflammation, metabolism, and cell stress or death ^28^. Peri-infarct microglia exhibited upregulation primarily in the immune response functional pathway (Fig. 3b-d), whereas neurons showed upregulation of cell stress / death pathways. The HK2 inhibitors attenuated not only the changes observed in peri-infarct microglia but also peri-infarct neurons, but had negligible effect on neuronal gene expression in the contralateral cortex (Suppl. Fig. 4c).

### HK2 inhibitors attenuate morphological markers of microglial activation and neuronal injury

Assessments of microglial morphology were performed 48 hours after stroke, at a time when this response is near maximal. Microglia in the peri-infarct-1 cortex exhibited the typical features of pro-inflammatory microglial activation: enlargement of the cell soma and retraction of cell processes (Fig. 4). Microglia in the peri-infarct-2 region displayed similar but less marked changes (Suppl. Fig. 6). The morphological activation in both regions was partially suppressed by the HK2 inhibitors.

**Figure 4.**
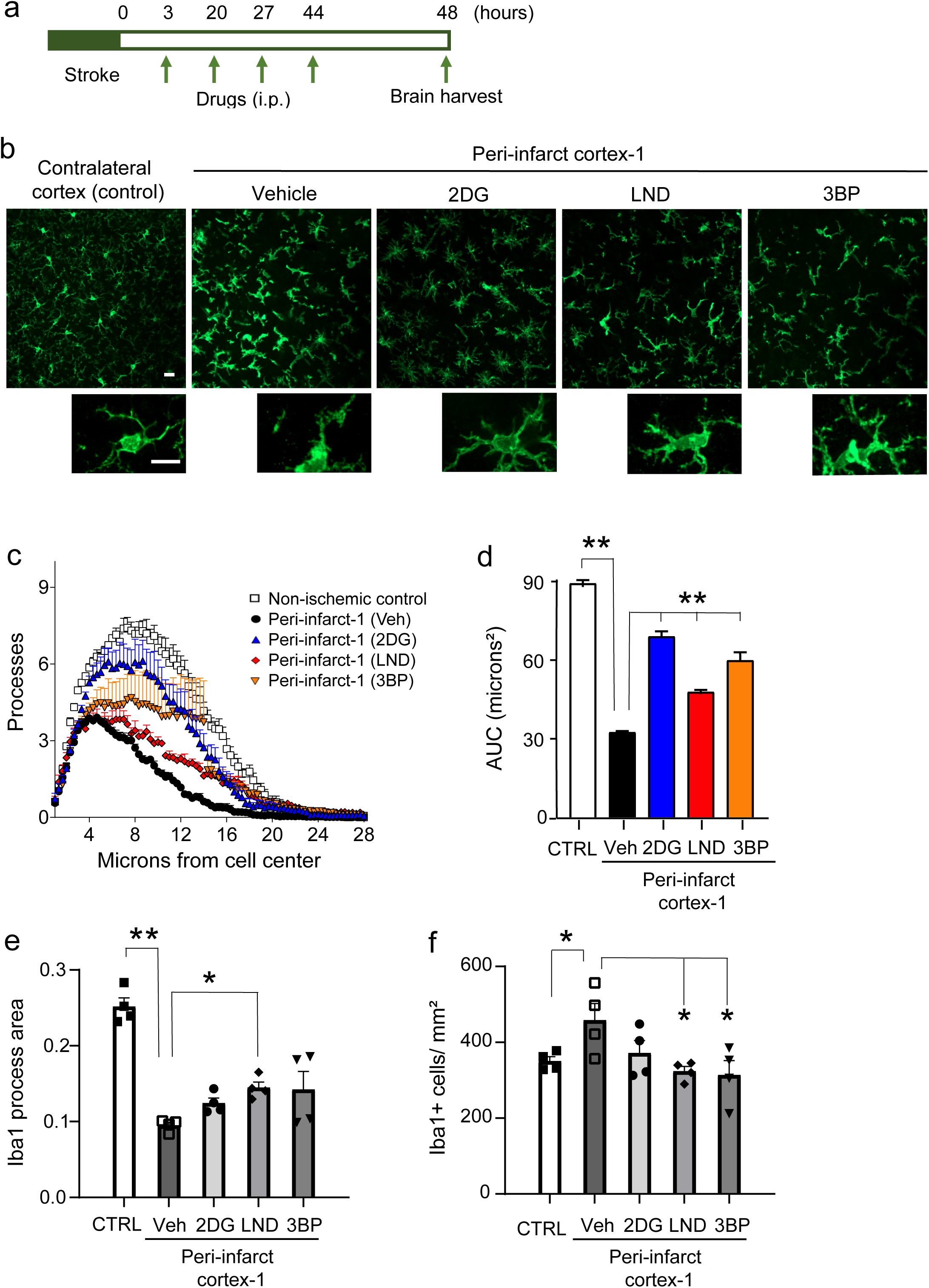
Effects of HK2 inhibitors on microglial morphology. **a.** Experimental design. **b**. Microglial morphology as shown by Iba1 immunolabeling. High-magnification images show cells with morphology representative of each treatment group. Scale bars = 10 µm. **c,d.** Sholl analysis shows the mean number of microglial processes at distances from the cell center. These are quantified as area under the curve. **e**. Mean microglial process area. **f**. Microglia density. n = 4; * p < 0.05, ** p < 0.01 by ANOVA and Dunnett’s test.

Effects of HK2 inhibitors on secondary neuronal injury were also examined at the 48-hour time point. There was no detectable loss of neuronal cell bodies in peri-infarct cortex, but there was a more than 50% reduction in neurite density (Fig. 5). This reduction is consistent with the particular sensitivity of axons and dendrites to inflammation-induced injury ^29, 30^ The neurite loss was reduced by the HK2 inhibitors, particularly LND and 3BP (Fig. 5). Inflammation-induced neurite loss results in part from the formation of cofilactin rods in response to local oxidative stress ^29^. Here, cofilactin rod formation and oxidative stress were evident as early as 6 hours after stroke (Fig. 6), with cofilactin rods identified as rod-like accumulations of cofilin-1, and oxidative stress indicated by nuclear DNA damage. All three HK2 inhibitors suppressed both neuronal oxidative stress and cofilactin rod formation, with LND and 3BP having larger effect sizes (Fig. 6).

**Figure 5.**
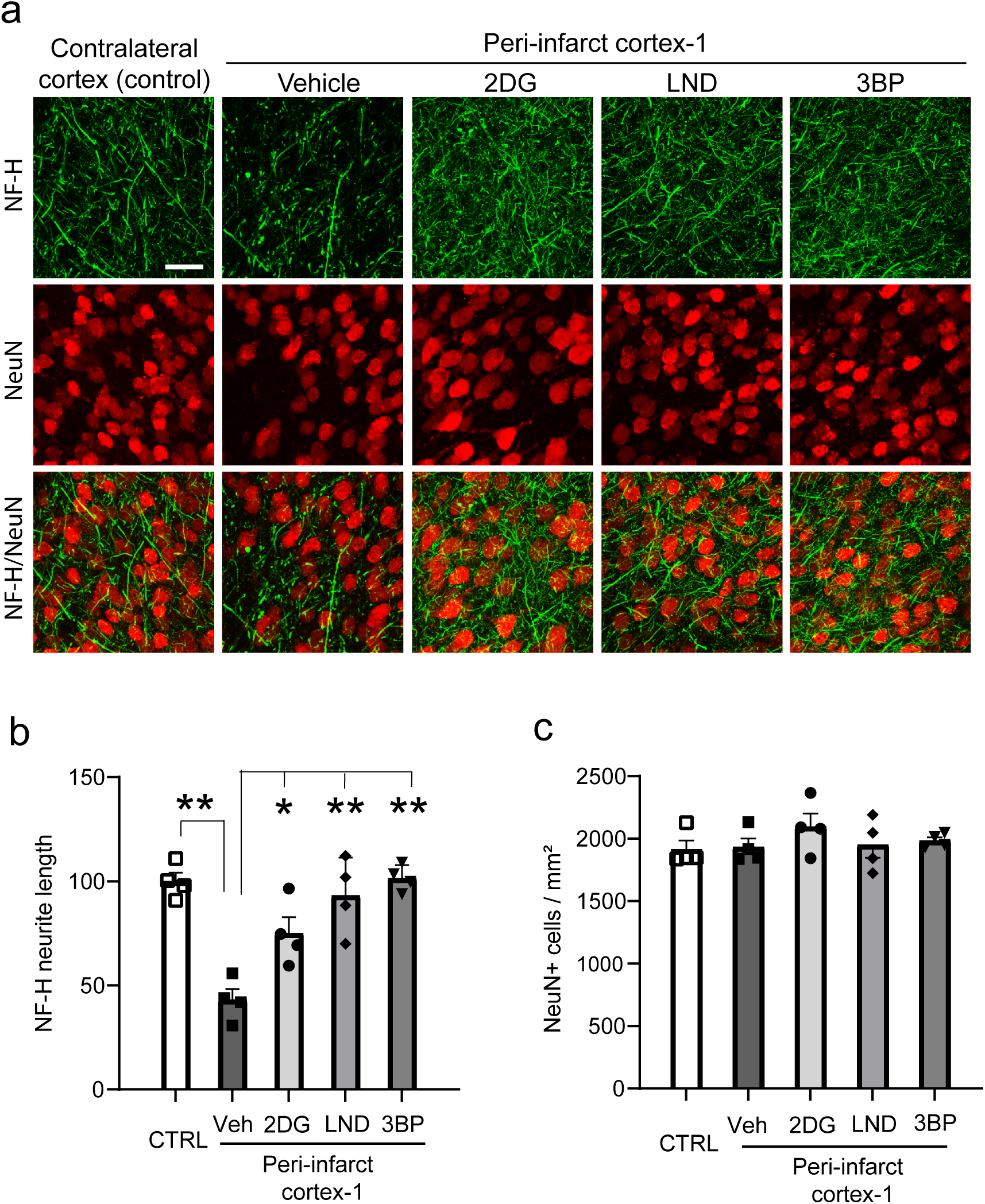
HK2 inhibitors administered after stroke reduce peri-infarct neurite loss. **a.** Neurites are identified by immunostaining for neurofilament-H (NF-H, green), and neuronal soma by NeuN (red). **b.** Neurite density expressed as total NF-H process length / neuron and normalized to the control (contralateral) cortex. **c**. There was no detectable loss of neuronal cell bodies in the peri-infarct cortex-1. * p < 0.05, ** p < 0.01 vs. vehicle by ANOVA and Dunnett’s test.

**Figure 6.**
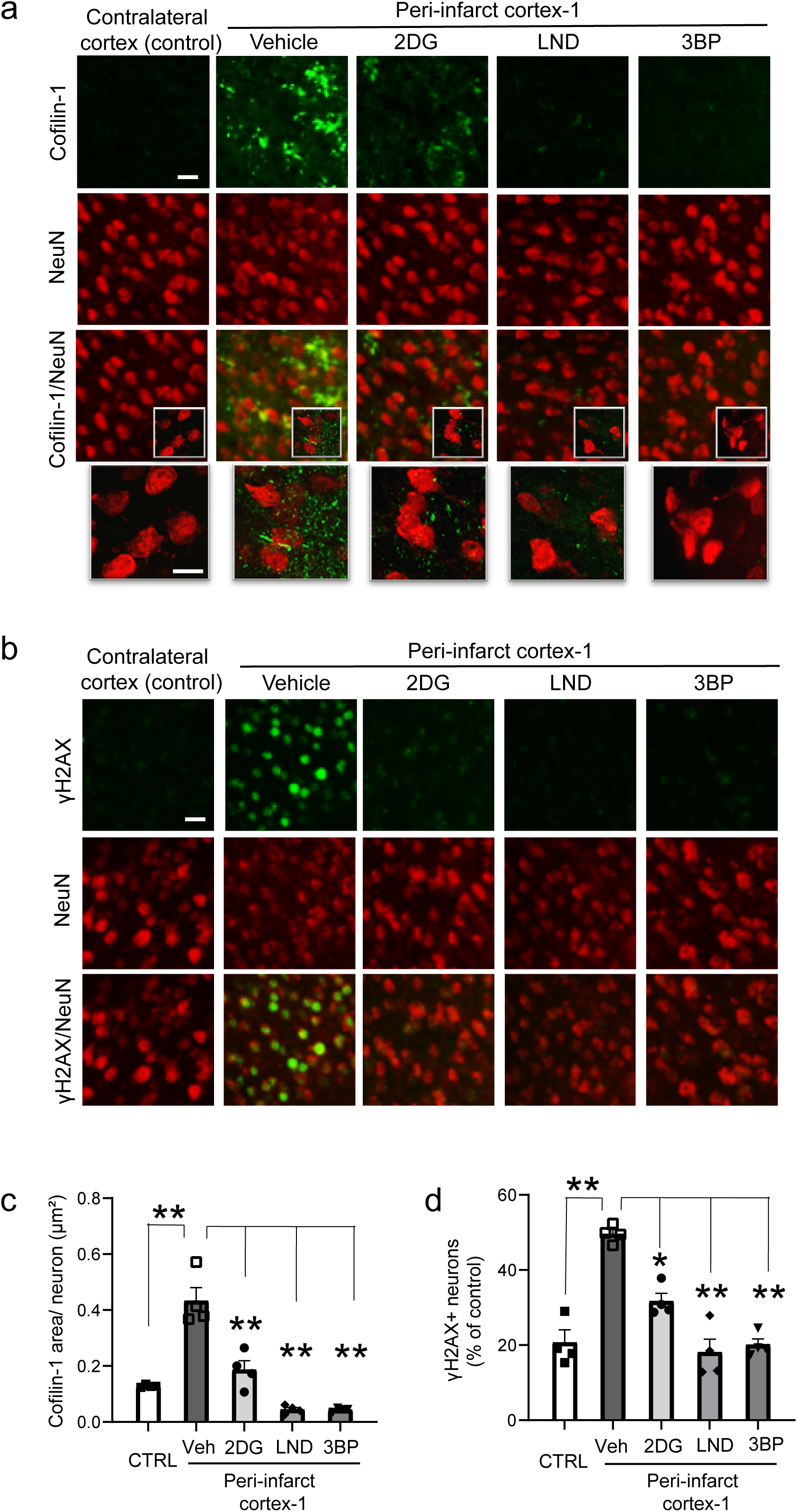
HK2 inhibitors suppress cofilactin rod formation and neuronal DNA damage in peri-infarct cortex. **a.** Formation of cofilactin rods (cofilin-1, green) in peri-infarct-1 cortex 6 hours after stroke. High magnification views show rod-like morphology of the cofilin-1 aggregates. Scale bars = 10 µm. **b.** DNA damage as identified by foci of γH2Ax formation (green) in neurons (NeuN, red) of peri-infarct-1 cortex at 6 hours after stroke. **c**. Quantification of cofilactin rod formation. **d**. Quantification of γH2Ax formation, expressed as number of cells with γH2Ax signal higher than the 80^th^ percentile of neurons in the control (contralateral) cortex. n = 4, * p < 0.05; ** p < 0.01 vs. vehicle by ANOVA and Dunnett’s test.

The effects of HK2 inhibitors on functional recovery were evaluated in mice treated with 3BP (or vehicle) initiated 3 hours after stroke, with dosing repeated twice daily for the subsequent 3 days (Fig. 7a). The mice were serially assessed the cylinder test, the rotating beam test, and the skilled reaching test over 21 days. The cylinder test showed increased motor asymmetry in both groups over the initial days after stroke followed by gradual recovery, with the 3BP-treated group having overall better (less asymmetric) performance. The rotating beam test, in which the rotation speed is periodically increased, revealed superior performance in the 3BP-treated mice after each increase. The skilled reaching task similarly showed superior performance in the 3BP-treated group sustained over the entire 21-day testing interval (Fig. 7b-d). Infarct size was measured in brains harvested at the end of the 21-day testing interval, and was slightly smaller in the 3BP-treated group (Fig. 7e,f).

**Figure 7.**
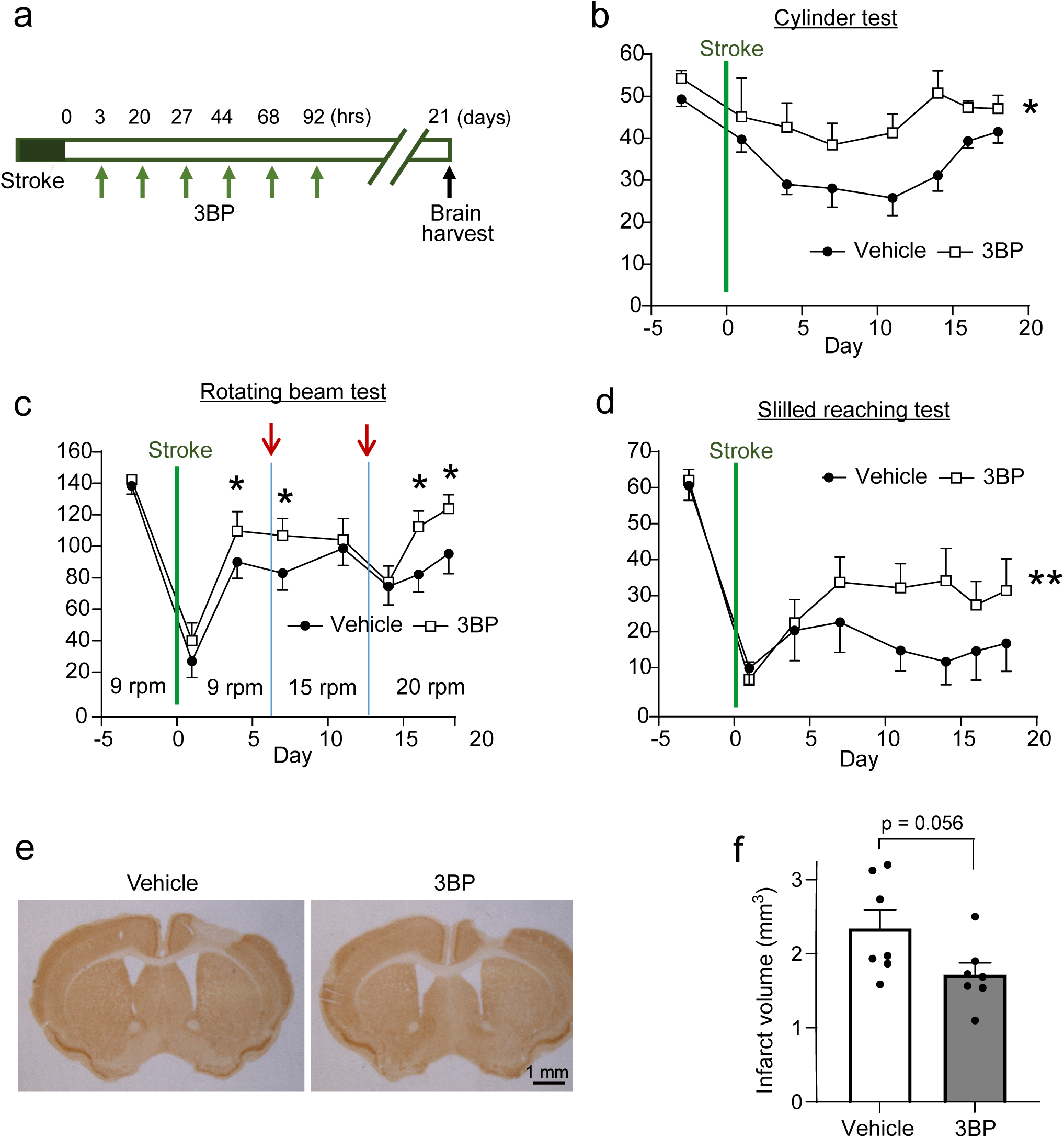
HK2 inhibitor administered after stroke reduce motor deficits and infarct size. **a.** Experimental design. **b**. Cylinder test. *p < 0.05 by repeated measures ANOVA over time (n = 7). **c**. Rotating beam test. Arrows denote days on which beam rotation speed was increased. * p < 0.05 by Student’s t-test on the designated days after speed increase (n = 15). **d**. Skilled reaching test. **p < 0.05 by repeated measures ANOVA over time (n = 7). **e.** Coronal sections of mouse brain immunostained for NeuN to identify the infarct. **f**. Brains harvested 21 days after stroke. Quantified infarct volumes in mice treated with 3BP or vehicle only following stroke. p = 0.05, n = 7.

## DISCUSSION

Our findings show that HK2 inhibitors can suppress the pro-inflammatory transcriptional response in peri-infarct microglia and reduce the cell stress/death transcriptional response in neighboring neurons. These transcriptional changes were associated with reduced peri-infarct microglial morphological activation, reduced neuronal oxidative stress and neurite loss, and improved functional outcome.

3BP, LND and 2DG all inhibit HK2 catalytic activity, but 2DG also inhibits HK1, and 3BP and lonidamine can have other off-target effects at higher doses ^16, 31, 32^. It was thus of interest to compare the effects of these agents to one another. 3BP and LND had nearly identical effects on microglial gene expression, both in both pattern and magnitude, and likewise had nearly identical effects on microglial morphology and neuronal injury. These findings are consistent with a shared mechanism of action. Additionally, the much smaller effects of these agents on the microglial and neuronal transcriptome in the un-injured cortex, where HK2 expression is not upregulated, suggests an absence of significant off-target effects. The non-selective HK inhibitor 2DG had a somewhat different pattern of effects on peri-infarct microglial and neuronal gene expression, and had smaller effects on microglial morphology changes, neuronal oxidative stress. and neurite damage. The smaller magnitude of the 2DG effects may reflect a lesser degree of HK2 inhibition at the dose administered. The 2DG dose was limited by mouse sedation, likely due to neuronal HK1 inhibition and resultant suppression of neuronal glycolysis^33, 34^.

The spatial transcriptomic approach permits simultaneous evaluation of gene expression in microglia and neighboring neurons. The functional pathway analyses showed that the pro-inflammatory microglial changes were associated with upregulation of neuronal stress/death pathways in both the peri-infact-1 and peri-infact-2 regions and that these changes in both microglia and neurons were lessened by the HK2-selective inhibitors. Given the near-absence of the HK2-selective inhibitors on neurons in undamaged cortex, together with the negligible HK2 gene expression detected in neurons, these results suggest that the HK2 inhibitor effects on neurons were indirect and secondary to their effects on microglia. The reduced neuronal oxidative stress in the HK2 inhibitor - treated peri-infarct cortex provides a potential mechanism for this interaction.

We used a random forest classification to identify the gene set most characteristic of peri-infarct microglia compared to contralateral microglia. Genes and their mechanisms in this set were found to overlap with the DAM gene set described in microglia associated with Alzheimer’s disease pathology ^24^, which exhibit innate immune activation and an increase in glycolytic metabolism. The upregulation of the DAM gene set in peri-infarct-1 microglia and suppression of this upregulation by HK2 inhibitors further suggests shared inflammatory and metabolic pathways between these microglial states ^5, 35, 36^.

The *in situ* transcriptomics approach enabled a simultaneous evaluation of transcriptional and morphological changes in microglia. In the peri-infarct-1 region, HK2 inhibitors ameliorated the upregulation of pro-inflammatory gene expression in microglia at 12 hours after stroke and correlated with the classical morphological changes observed at 48 hours. Changes in the more distant peri-infarct-2 microglia were much smaller in magnitude, and the changes in microglial morphology were likewise less pronounced. These observations confirm a correlation between the gene expression and morphology changes, but with the caveat that these gene expression assessments were intentionally performed at a time point preceding the histological evaluations, assuming a time lag between the two. Moreover, it is uncertain what transcriptional changes drive microglial morphology changes, or even if these are transcriptionally regulated.

The neurites (axons and dendrites) by which neurons communicate with one another have recently been shown to be particularly vulnerable to inflammation-induced injury through a process involving oxidative stress and formation of intra-neurite cofilin-1 / actin aggregates (cofilactin rods). Our results here confirm oxidative stress, cofilactin rod formation, and neurite loss in peri-infarct cortex, all of which were attenuated by the HK2 inhibitors. The infarct size was also slightly smaller in the 3BP-treated mice, despite the 3 - hour interval between stroke and first drug administration, a result similar to that reported using delayed administration of fingolimod after permanent ischemia ^37^. These findings, along with the attenuated upregulation of cytokine and protease genes in the mice treated with the HK2 inhibitors, provide plausible mechanisms by which functional outcome was improved by 3BP.

The *in situ* transcriptomics approach employed here eliminates artifacts induced by gene expression changes during tissue dissociation and sorting during standard cell isolation methods ^38^ and allowed precise exclusion of microglia that are either remote from the infarct or had migrated into the infarct itself and thus unlikely to affect peri-infarct neuronal injury. Prior studies of microglial gene expression after stroke have used a variety of stroke models, tissue sampling areas, and methods for assessing gene expression ^39, 40, 41, 42, 43, 44, 45, 46, 47, 48, 49, 50^. These methodological differences complicate comparisons between studies, but our results are in general accord with the previous reports identifying upregulation of cytokine, chemokine, and protease-encoding genes in microglia. Prior studies using spatial transcriptomics in mouse models of stroke also highlight spatial heterogeneity in microglial responses ^46, 47, 48, 50, 51^.

Our findings contribute to the existing literature on glycolytic inhibition and the innate immune response. Ketogenic diet, which limits glucose flux through glycolysis, can suppress the innate immune response in a variety of settings, including stroke ^52, 53, 54^. It was shown in the 1950s that injury-induced inflammation can be robustly suppressed by 2DG ^55^, and 2DG was subsequently found to have anti-inflammatory effects in a variety of conditions including stroke ^23, 56, 57^. 2DG is not a feasible therapeutic after stroke because the resulting suppression of HK1 activity in neurons and other cell types could exacerbate energy compromise in damaged or marginally perfused tissue. Salutary effects of selective HK2 inhibition have been reported in stroke ^10, 16^, though not using the clinically relevant, delayed administration or with the long-term behavioral endpoints assessments used here.

Several mechanisms have been identified by which glycolytic inhibition may affect the innate immune response. A reduced glycolytic flux reduces cytosolic NADH levels, which in turn suppresses the action of the master inflammatory transcription factor NFκB by promoting dimerization (inactivation) of the NADH –sensitive co-repressor CtBP ^58^. In addition, non-transcriptional effects may be mediated by reduced capacity to generate the reactive oxygen species superoxide and nitric oxide, both of which need glucose as an obligate precursor ^59, 60, 61, 62, 63, 64, 65^. Recent reports also identify additional, non-glycolytic actions specifically of HK2 that can influence immune responses. HK2 untethered from mitochondria can promote inflammasome assembly and thus IL1β production ^66^. Untethered HK2 can also phosphorylate IκBα to cause IκBα proteolysis and resultant nuclear translocation of NFκB ^67^, but the effect of Hk2 inhibitors on these processes is not known. Our results demonstrated that the upregulation of HK2 gene expression in peri-infarct microglia was suppressed by HK2 inhibitors, suggesting that ongoing HK2 activity was required for maintaining the elevated HK2 expression. The interruption of this feed-forward loop may explain why HK2 inhibitors are particularly effective in suppressing the innate immune response.

Unexpectedly, the astrocyte-specific gene GFAP was among the genes robustly upregulated in both microglia and neurons of peri-infarct in the unfiltered transcriptome data set. A similar result was reported by ^68^ also using a spatial transcriptomic approach. GFAP is one of several genes that are highly upregulated in reactive astrocytes ^69, 70^. Given that GFAP mRNA is transported through distal astrocyte processes ^71, 72, 73^, we suggest that the GFAP signal identified in peri-infarct microglia and neurons arises from mRNA in astrocyte processes traversing microglial and neuronal somas identified for probe photocleavage. This idea is supported by the increased detection, in both peri-infarct-1 neurons and microglia. of other mRNA transcripts that are found in astrocyte processes together with the lack of any increase in the nucleus-specific astrocyte transcripts SOX9, NFIA or S100B ^71^. We therefore excluded the genes known to be expressed in astrocyte processes from the subsequent analysis of the microglial and neuronal transcriptome data ^71^ (Supplemental Data File 13).

A potential limitation to these studies is the use of contralateral, uninjured cortex as control rather than cortex from uninjured mice. This approach permits each mouse to serve as its own control; however, and even though the photothrombotic stroke model requires no craniotomy or neck incision, it remains possible that the stroke on one hemisphere may have had effects on gene expression or histological markers on the contralateral side. An additional limitation is that contributions of astrocytes and infiltrating neutrophils were not directly assessed in this study, and that only singe time-points were used for the transcriptomic and histological evaluations.

## METHODS AND MATERIALS

### Animals

Studies were approved by the animal studies committees at the San Francisco Veterans Affairs Medical Center, and were performed in accordance with the National Institutes of Health Guide for the Care and Use of Laboratory Animals. Data were acquired and reported in accordance with the ARRIVE 2.0 guidelines ^74^. C57BL/6 mice were obtained from the Jackson Laboratories. Each experiment included equal numbers of male and female mice, age 3 - 5 months old. There were no premature animal deaths during the study. Mice were arbitrarily assigned to the various treatment conditions, with gender balance and approximately equal ages in all groups.

### Stroke induction

Mice were anesthetized with 2% isoflurane in 70% N_2_O/ balance O_2_ delivered through a ventilated nose cone. Rectal temperature was maintained at 37.0 ± 0.5 °C with a homeothermic blanket throughout the surgical procedure. A photothrombotic infarct was induced by the Rose Bengal technique ^30, 75^. For studies involving functional recovery, the infarct was placed on the motor cortex contralateral to the dominant paw, as assessed during acclimation to the skilled reaching task. After exposing the skull by skin incision, an isosceles right triangle-shaped adaptor (3 mm base x 3 mm height) was centered over the primary motor cortex (1.0 mm anterior, and 1.5 mm lateral to bregma) and connected to a white light source (KL 1500 LCD, SCHOTT North America Inc., Southbridge, MA) with a 2 mm diameter fiber optic cable. Rose Bengal (Sigma-Aldrich, St Louis, MO; 20 mg/kg) dissolved in saline was infused into the retro-orbital sinus for 30 seconds, and light was transmitted through the fiber optic cable for 15 minutes. Sham ischemia animals were injected with saline only but were otherwise treated identically. The incisions were sutured, bupivacaine (6 mg/kg) and buprenorphine SR (0.1 mg/kg) were administered subcutaneously, and the mouse was moved to a warm recovery chamber until awake and ambulatory. Vehicle solution (10% DMSO in saline), 2-deoxyglucose (2DG, 250 mg/kg), lonidamine (LND, 50 mg/kg) and 3-bromopyruvate (3BP, 5 mg/kg) were injected intraperitoneally (i.p.) at the indicated time points.

### Immunohistochemistry

Anesthetized mice were perfused with cold saline (0.9% NaCl) followed by a 4% solution of paraformaldehyde in phosphate-buffered saline, pH 7.4 (PFA). Brains were removed and post-fixed with PFA for 24 hours, then immersed for another 24 hours in 20% sucrose for cryoprotection. The brains were then frozen and 40 µm coronal sections were prepared with a cryostat. The fixed brain sections were pre-incubated in a blocking buffer (2% donkey serum, 0.3% Triton X-100 and 0.1% bovine serum albumin in 0.1 M phosphate buffer) at room temperature for 1 hour and then incubated with the primary antibodies overnight at 4 °C. The antibody sources and dilutions used are listed in Supplemental Table 1. After washing, antibody binding was detected using fluorescent secondary antibodies listed in Supplemental Table 1.

Stained sections were mounted on glass slides in a DAPI-containing anti-fade mounting medium (Vector laboratories, Burlingame, CA). For detection of cofilactin rods, the fixed sections were incubated with 100% methanol at -20 °C for 15 minutes and incubated with anti-cofilin-1 antibody in blocking buffer lacking detergent. Immunolabeling for NeuN was performed in a second step using standard, detergent-containing blocking buffer.

### Spatial gene expression

Paraffin-embedded brains were cut into 5-µm coronal slices, mounted on glass slides, and stored at -20 °C with desiccant. The sections were deparaffinization and treated with Tris-EDTA (pH 9.0) for antigen retrieval and then incubated overnight with whole-mouse transcriptome RNA probes linked with photo-cleavable DNA tags ^76^. The washed sections were then incubated with the DNA-binding fluorophore SYTO83 as a nuclear marker and with cell-type specific antibodies (anti-NeuN and anti-Iba1) visualized by subsequent incubation with secondary antibodies conjugated with green and far-red fluorophores. The DNA probes were then cleaved from cell types and regions of interest by UV light illumination (Fig. 1a,b). Slides were loaded onto a GeoMx Digital Spatial Profiler (Bruker Spatial Biology) and scanned to generate digital images of the fluorescent markers. Three regions of interest were selected: (1) a parallelogram bordering on the lateral edge of the infarct margin (as identified by loss of NeuN-positive neurons) and extending 500 μm laterally, termed here as “peri-infarct-1 cortex”; (2) a second parallelogram extending and additional 500 μm laterally, termed “peri-infarct-2 cortex”; and (3) the homologous contralateral (uninjured) cortex, termed “contralateral cortex” (Fig. 1a,b). Within each region of interest, the photocleavable DNA tags connected to the transcriptome probes were liberated with 1 μm columns of UV light were sequentially targeted to the nuclei of microglia and neurons, as identified by the immunolabeling. The DNA tags contained a sequence identifier, a unique molecular identifier, and primer binding sites. Following each UV illumination the cleaved tags were collected by microcapillary aspiration into a 96-well plate. The collected oligonucleotides were amplified by PCR, purified with AMPure XP beads, and then quality assessed using QuBit. The amplified libraries were then sequenced using a NovaSeq 6000 sequencer (Illumina).

### Histology image analysis

The regions of interest used for histology image analysis were defined in the same way as for spatial gene expression studies. Three sections per mouse, spaced 240 µm apart, were used for each assessment. Image quantification was performed by observers blinded to the mouse treatment conditions, as described ^29^. For γH2Ax, the mean integrated fluorescence density was measured in the neuronal nuclei (as identified by NeuN immunolabeling) and expressed as the percent of neurons in which the signal intensity exceeded the 80^th^ percentile value of neurons measured in homologous contralateral (uninjured) region ^77^. Neurite length measurements were made after thresholding the NF-H images using the ImageJ Otsu function. The total NF-H neurite length in each image was quantified by summing the neurite lengths after skeletonizing the images in ImageJ and then normalized to the number of neuronal cell bodies per image as assessed by NeuN immunostaining ^29^. For analysis of cofilactin rod formation, the images were thresholded using the Fiji Triangle function, and the area of cofilin-1 immunofluorescence was summed and normalized to the number of neurons per image.

For Sholl analysis of microglial arborization ^78^, Z stacks of Iba immunostained images were compressed to form maximum intensity projections and thresholded using the “Make Binary” function in ImageJ. Sholl analysis was conducted on individual cells using the Sholl plugin (ImageJ) with start radius = 0 pixels and step size = 1 pixel. Every fourth cell was analyzed in each image, excluding cells whose processes were cut off by the image edge. Data are displayed as the number of intersections formed by microglial processes at each radius from the cell center and the area under the curve (AUC) for each mouse. A separate analysis of total Iba1 process area per image (excluding the cell body area) was performed using the ImageJ particle size function.

### Infarct volume

Brain infarct volume was assessed in mice used in the behavioral studies, at 21 days after ischemia. Twelve brain sections spaced 240 µm apart, spanning the entire infarct region, were immunostained with anti-NeuN and visualized by the diaminobenzidine method ^79^. The area of neuronal loss in each section was calculated in Image J software, and the infarct volume in each brain was calculated summing these for each animal and multiplying by the 2.88 mm spanned by the 12 sections.

### Behavioral studies

Mice were evaluated with three tests to assess locomotor asymmetry and motor dexterity: the cylinder test, rotating beam test, and skilled reaching test. The mice were acclimated to handling over 2-weeks before stroke, during which time they were also acclimated to the rotating beam and the skilled reaching tasks. Post-stroke testing was performed on days 1, 4, 7, 10, 14, 15, and 17 days after stroke. All observers were blinded to the mouse treatment conditions. For the cylinder test, a mouse was placed in a beaker (12 cm height x 17 cm diameter) for 10 minutes while being video-recorded from above. ^80^. The beaker was washed with 10% bleach in between mice. The videos were reviewed to determine which forelimb touched the beaker surface during rearing behavior. Touches were counted when they occurred immediately after a mouse rearing, and no score was assigned if the mouse touched the beaker wall with both forepaws simultaneously. Results were expressed as the percentage of touches made by the impaired forelimb (contralateral to infarct) relative to the total touches recorded [(touches by contralateral forelimb) / (touches by either forelimb) x 100].

The Wishaw skilled reaching task was performed using a custom-made automated box as previously described for rats ^81, 82^, size-modified for use in mice. The mice were trained to reach through a 0.5 cm slit for a 14 mg rodent diet pellet (Bio-Serv, # F0071) before induction of stroke. Paw preference was noted during initial training sessions, and a reaching slit was placed to facilitate the use of the preferred paw. Mice were fasted overnight before the training sessions and fed 10% of their body weight after the training (minus the amount eaten during training). Mice were considered adequately trained when they could successfully grab and eat a pellet in over 30 of 60 consecutive trials. Approximately 60% of the mice achieved this metric and those that did not were excluded from the studies. Stroke was induced in the hemisphere contralateral to the dominant paw. During post-stroke testing, mice were given 60 opportunities to grab a pellet, and the percentage of successful grabs (pellets eaten) was scored.

The rotating beam test ^83^ requires mice to traverse the length of a rotating 145 cm long / 1 cm diameter beam. The task is scored by the distance the mouse covers before falling off onto a safety net below. The mice were habituated to the rotating beam, with rotation speed increased from 0 rpm to 9 rpm. The mice completed 3 repetitions on the beam on each day of training. Mice displaying right paw preference were trained on a clockwise rotating beam and those with left paw preference on a counter-clockwise rotating beam. The beam was wiped clean with bleach between mice. Mice were considered adequately habituated when they could traverse the full 145 cm at a 9 rpm rotation speed on 3 consecutive trials. After stroke, the beam rotation speed was set at 9 rpm on days 1 and 4, at 15 rpm on days 7 and 10, and then at 20 rpm on days 14, 15, and 17. Each testing session consisted of 3 repetitions on the beam. The mean distance traveled was recorded as the score obtained for each testing day.

### Bioinformatics and Statistics

Sequencing reads were aligned to mouse genome (MGSCv37) for production of raw counts. The raw gene counts were filtered for lowly expressed genes (<30 CPM averaged across samples). We additionally filtered 214 genes known to be expressed in astrocyte processes ^71, 72, 73^, because the spatial transcriptomics method cannot exclude RNA from astrocyte processes traversing the neuronal and microglial nuclei of interest. These are listed in Supplemental Data File 13. The remaining gene counts were normalized with the variance stabilizing transformation algorithm, and differential gene expression was performed in R with DESeq2 ^84^. Random forest classification and variable importance analysis for activated microglia gene signature generation were performed in R with nestedCV using nested cross-validation of the normalized microglial gene expression matrix of vehicle treated mice as the input ^85^. Gene set variance analysis was performed using gene ontology (GO): Biological pathways annotations and the GSVA package^28^. Gene expression plots, heatmaps, and volcano plots were generated with ggplot2, pheatmap, and EnhancedVolcano, respectively ^86, 87, 88^.

Histology data are expressed as means ± s.e.m., with the “n” of each experiment defined as the number of mice or, for cell cultures, the number of independent experiments. Results were analyzed by unpaired t-test when only two groups were compared, by ANOVA with Dunnett’s test for comparisons against a common reference group.

## AUTHOR CONTRIBUTIONS

R.A.S, SJ.W. and G.U. conceived the experiments. SJ.W., G.U., N.M., C-L.T., D.O., S.A., K.K., E.M., R.S, K.A., and W.C. performed the experiments, K.A., N.S. SJ.W., G.U., and R.A.S. analyzed the results, and SJ.W, G.U., N.S. and R.A.S. wrote the manuscript. All authors reviewed the manuscript.

