## Supplemental Table and Figures for "Transcriptional and neuroprotective effects of hexokinase-2 inhibitors administered after stroke"

**Supplemental Table 1: Table of antibody sources and dilutions used.**

| Antibody | Source | ID | Host | Dilution |
|---|---|---|---|---|
| Primary Antibodies | | | | |
| Cofilin | Millipore, Temecula, CA, USA | 51755 | Rabbit | 1:300 |
| γH2Ax | Cell Signaling, Danvers, MA, USA | 9718S | Rabbit | 1:500 |
| Iba1 | Wako, Richmond, VA, USA | 019-19741 | Rabbit | 1:1000 |
| NeuN | Millipore, Temecula, CA, USA | ABN78 | Rabbit | 1:1000 |
| NeuN | Synaptic systems, Temecula, CA, USA | 266004 | GuineaPig | 1:500 |
| Neurofilament-H | BioLegend, San Diego, CA, USA | 801601 | Mouse | 1:500 |
| Secondary Antibodies | | | | |
| Cofilin | Vector Lab, Burlingame, CA, USA | BA-1000 | Goat | 1:250 |
| anti-guinea pig IgG 594 | Jackson Laboratory, Bar Harbor, ME, USA | 106-585-003 | Goat | 1:1000 |
| anti-guinea pig IgG 647 | Thermo Fisher Scientific, Waltham, MA, USA | A21450 | Goat | 1:1000 |
| anti-mouse IgG 488 | Thermo Fisher Scientific, Waltham, MA, USA | A21202 | Donkey | 1:1000 |
| anti-rabbit IgG 488 | Thermo Fisher Scientific, Waltham, MA, USA | A21206 | Donkey | 1:1000 |

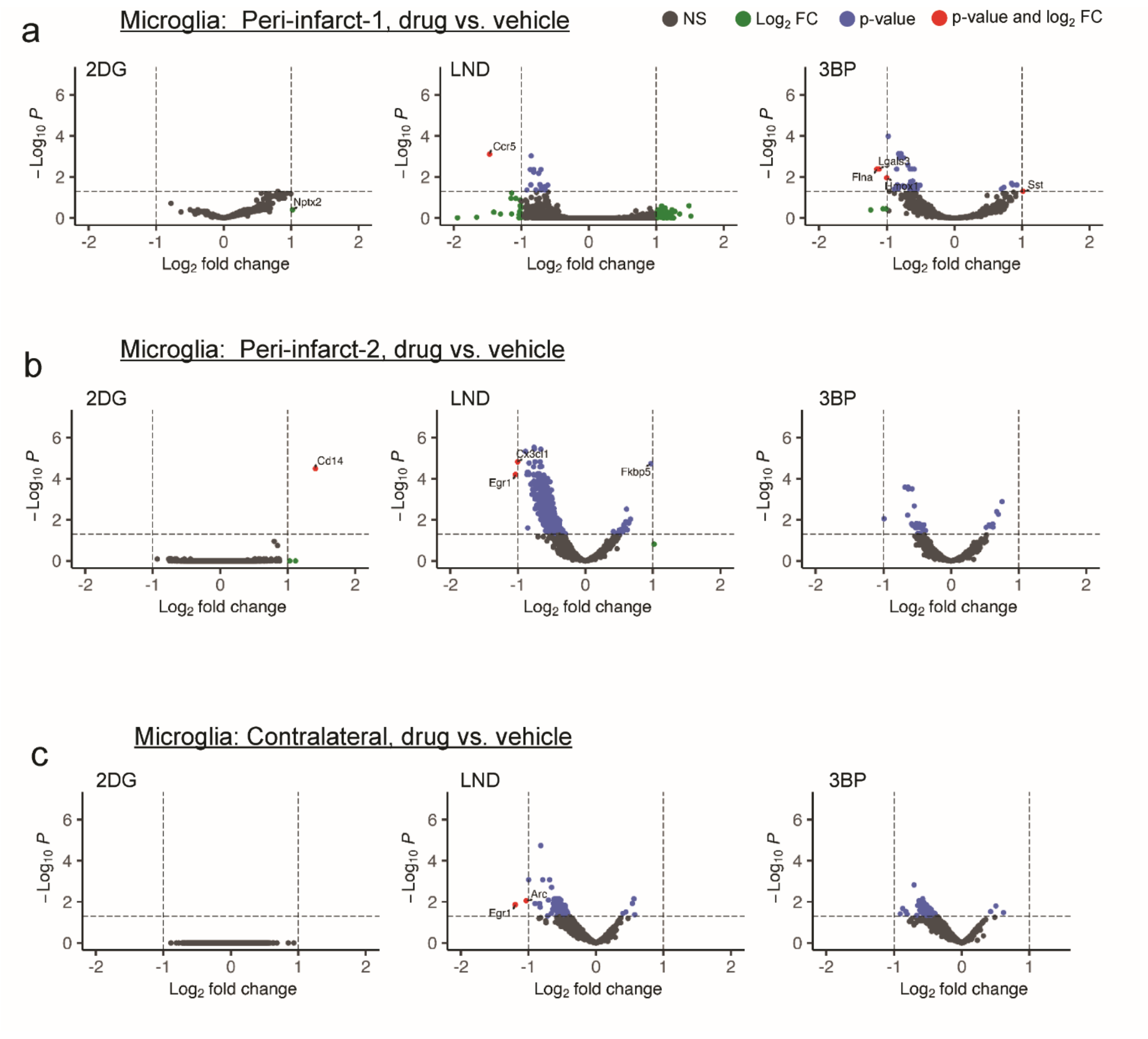


**Supplemental Figure 1. Microglial gene expression displayed as a comparison of each HK2 inhibitor vs. vehicle.** Data are from the (**a**) peri-infarct-1 cortex, (**b**) peri-infact-2 cortex, and (**c**) contralateral cortex. These data correspond to Fig. 2. Genes with log$_2$ expression fold change (FC) > 1 and p < 0.05 are denoted by red dots; log$_2$ FC > 1 and p > 0.05 by green dots; log$_2$ FC < 1 and p < 0.05 denoted by blue dots, and not significant (NS) by black dots.

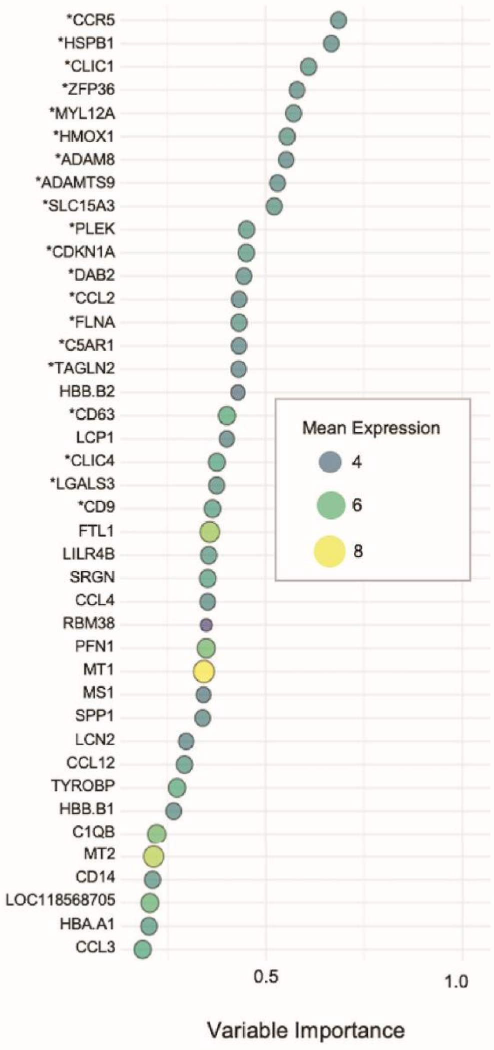


**Supplemental Figure 2. Gene set defining ischemia-activated microglia**

A random forest classification was used to identify a gene set in which expression changes are most characteristic of the changes observed in peri-infarct-1 microglia relative to contralateral microglia. Top scoring genes that were also highly expressed in microglia (denoted by *) comprised the activated microglial signature gene set displayed in Fig. 2. The size and color of the circles indicate the normalized gene expression levels.

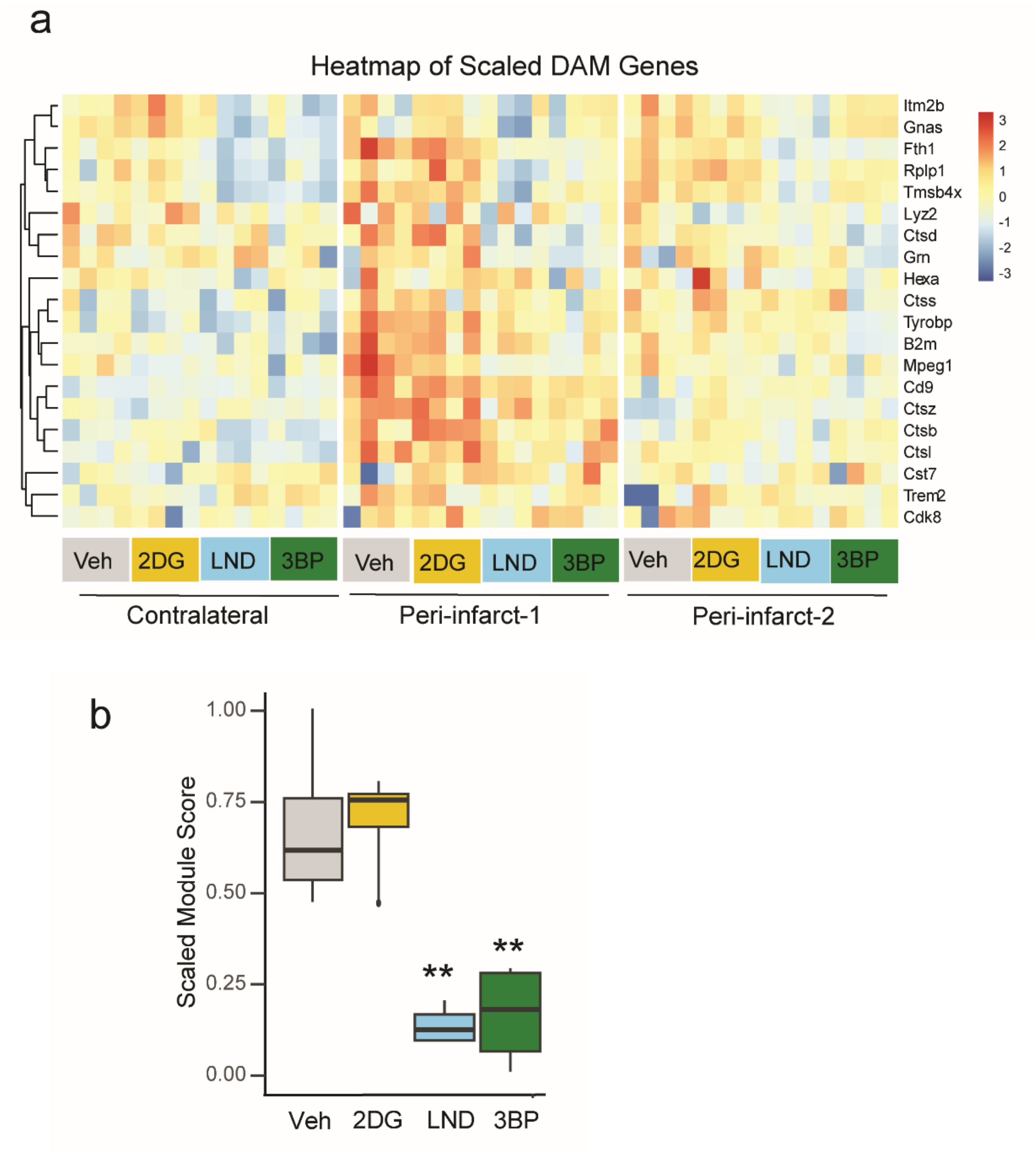


**Supplemental Figure 3. Microglial expression of neurodegenerative disease-associated genes**

**a**. Heatmap showing z-scaled expression of genes previously identified as upregulated in neurodegenerative disease-associated microglia (DAM) (Keren-Shaull et al., 2017). Data are shown for each treatment condition and region of interest. n = 4. **b.** Box plot showing effects of drug treatment on the normalized composite expression of the upregulated DAM gene set in peri-infarct-1 microglia. ** $p < 0.01$ vs vehicle by ANOVA with Dunnetts test.

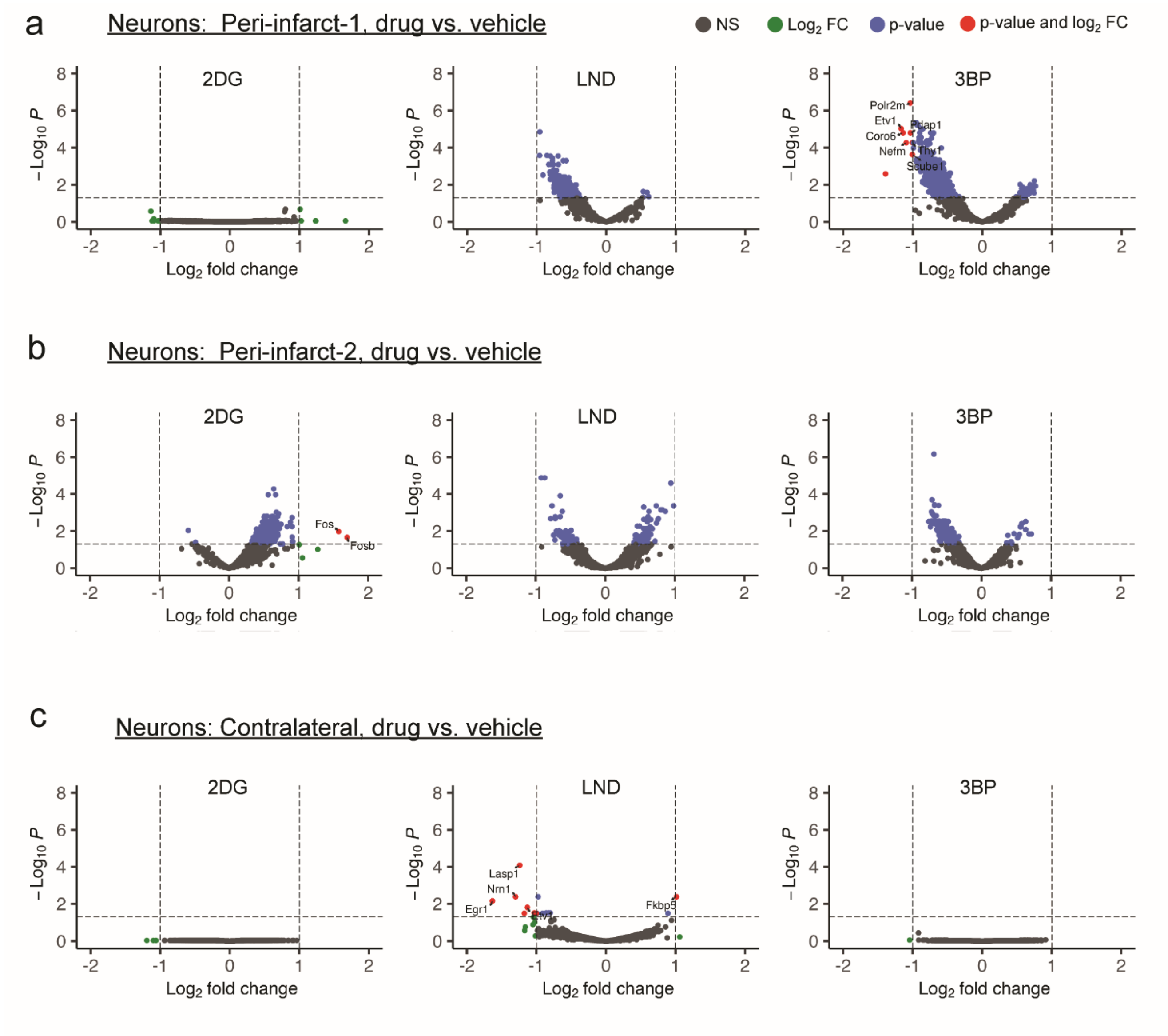


**Supplemental Figure 4. Effects of HK2 inhibitors on peri-infarct neuronal gene expression**

Data are from the (**a**) peri-infarct-1 cortex, (**b**) peri-infact-2 cortex, or (**c**) contralateral cortex. Genes with $\log_2$ expression fold change (FC) > 1 and $p < 0.05$ are denoted by red dots; $\log_2$ FC > 1 and $p > 0.05$ by green dots; $\log_2$ FC < 1 and $p < 0.05$ denoted by blue dots, and not significant (NS) by black dots.

**Immune Response**

1. Natural killer cell activation
2. Regulation of macrophae activation
3. TNF superfamily cytokine production
4. Macrophage chemotaxis
5. Macrophage migration
6. Monocyte chemotaxis
7. Activation of immune response
8. Cytokine production

**Metabolic**

1. ATP metabolic processes
2. Glucose-6-phosphate metabolic processes
3. Glutamate metabolic processes
4. Glutathione metabolic processes
5. Cellular homeostasis
6. Oxidative phopsphorylation
7. Regulation of mitochondrial membrane permeability
8. Superoxide metabolic processes

**Cell Stress/Death**

1. Apoptotic mitochondrial changes
2. Cell death in response to oxidative stress
3. Apoptotic signaling pathway
4. Cellular response to ROS
5. Cellular response to unfolded proteins
6. Neuronal death
7. Neuronal death in response to oxidative stress
8. Endoplasmic reticulum unfolded protein response

**Supplemental Figure 5. Gene functional pathways corresponding to Figure 3b.**

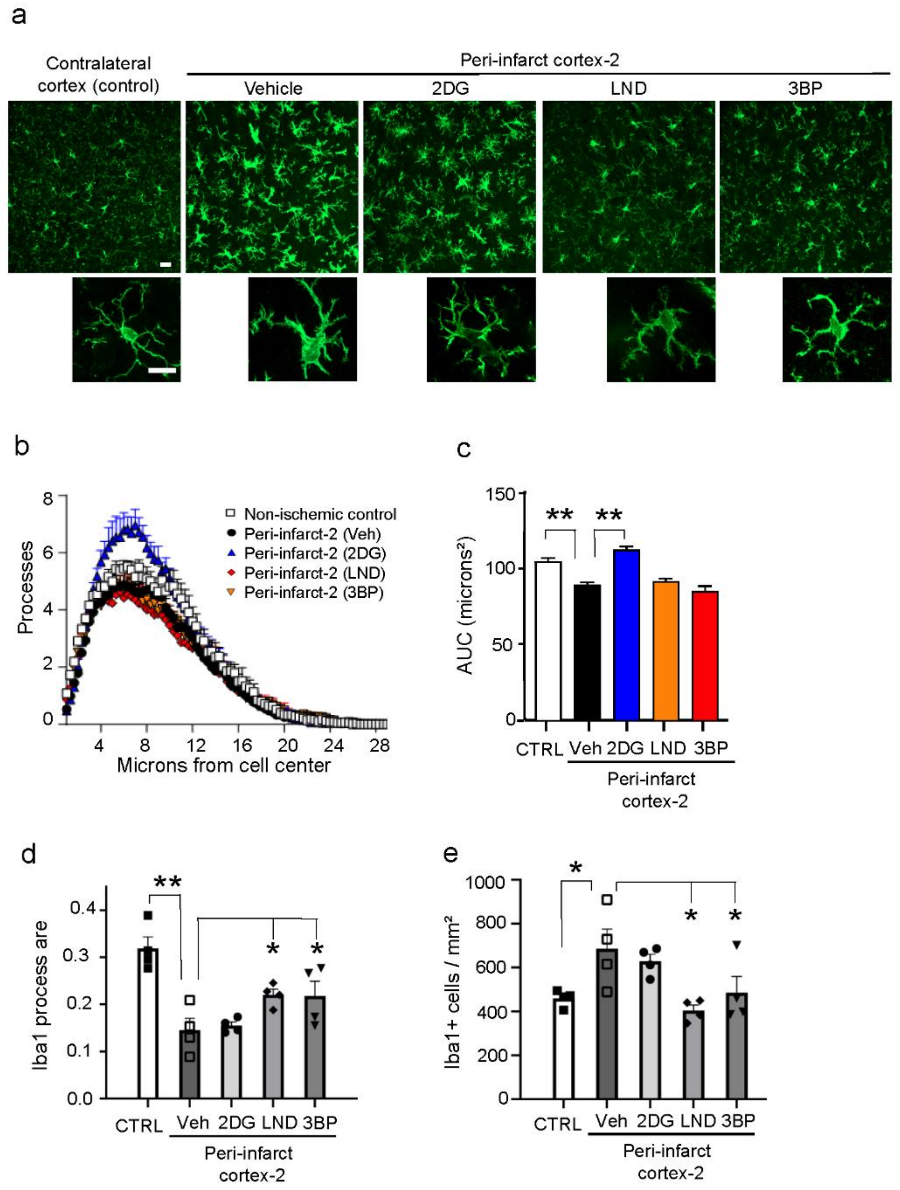


**Supplemental Figure 6   Microglia morphology in peri-infarct-2 cortex**

**a**. Microglial morphology as shown by Iba-1 immunolabeling. High magnification images show cells with morphology representative of each treatment group. Scale bars = 10 µm.  **b,c** Sholl analysis showing the mean number of microglial processes at distances from the cell center. This is quantified as area under the curve. **d**. Mean microglial process area. **e**. Microglia density. n = 4; * $p < 0.05$, ** $p < 0.01$ by ANOVA and Dunnett's test.
